# Microglial states determine lesion dynamics in multiple sclerosis

**DOI:** 10.1101/2024.10.25.620251

**Authors:** Aletta M.R. van den Bosch, Jia Hui Khoo, Zhigang Lu, Han Liang, Dennis Wever, Li Pu, Bart J.L. Eggen, Mathias Uhlén, Joost Smolders, Jörg Hamann, Zhouchun Shang, Jan Mulder, Inge Huitinga

## Abstract

Multiple sclerosis (MS) is a neuroinflammatory disease of the central nervous system, characterized by demyelinating lesions ^1^. Lesion expansion contributes to progression and increased disability, while remyelination can recover neurological deficits. However, mechanisms driving lesion dynamics are largely unclear, hindering the development of effective therapeutics. We propose that distinct states of microglia are involved in lesion expansion and remyelination ^2,3^. Using Stereo-seq, an RNA capture based high-resolution spatial transcriptomics technology with single-cell resolution, on post-mortem human brain tissue, we compared mixed active/inactive lesions with lipid-laden foamy microglia with lesions containing ramified microglia. We identified distinct cellular and molecular mechanisms underlying lesion activity and remyelination, linked to microglia phenotypes and states. Lesions with foamy microglia were characterized by elevated immune activation, increased lymphocyte densities, upregulated immunoglobulin production (*IGHG1*, *IGHG3*), increased complement system activity, indication of iron dysregulation (*FTL*, *FTH1*), and increased demyelination. In contrast, lesions with ramified microglia exhibited gene expression profiles indicative of myelin stability (*ABCA2*, *QKI*) and neuro-axonal protection, fostering an environment conducive to repair and remyelination. Our findings highlight the role of microglial states in lesion expansion and repair in MS and offer promising avenues for the development of therapeutic approaches aimed at preventing MS disability progression.

## Main text

Multiple sclerosis (MS) is the most prevalent neuro-inflammatory disorder among young adults, affecting more than 2.8 million people globally ^4^. Clinically, MS is characterized by recurrent episodes of partially reversible neurological dysfunction alongside a progressive accumulation of chronic neurological disability ^1^. Pathologically, MS is defined by focal areas of demyelination, known as lesion, distributed throughout the central nervous system (CNS). However, the disease course remains unpredictable, largely due to spatial and temporal pathological heterogeneity ^5^.

Ongoing lesion activity and failed remyelination are thought to contribute to disease progression, yet the molecular mechanisms underlying this process are not well understood. MS lesion types are classified based on the degree of de- or re-myelination and the accumulation of activated microglia in various states ^3^. Among these lesions, mixed active/inactive (mixed) lesions – characterized by a completely demyelinated core and a border of activated microglia – are particularly versatile. These lesions exhibit ongoing demyelinating activity while possibly retaining the potential to either remyelinate or to transit into inactive sclerotic scars ^3^. Despite their importance, the microglial states and the cellular and molecular mechanisms driving these processes could not be fully characterized due to technical limitations.

In this study, we employed Stereo-seq ^6^, a high-resolution, capture-based, unbiased spatial transcriptomics technology, with chips spanning up to 2 cm x 3 cm, with the aim to identify the cellular and molecular basis of MS lesion evolution of lesions by comparing lesions characterized by distinct microglial states. Microglia are highly dynamic cells. In the resting state, microglia have a ramified morphology, and after phagocytosis of lipids they can become foamy ^7^. We previously reported a correlation between lesions with foamy microglia and rapid disability accumulation ^8^, which may in part be explained by increased acute axonal damage and axonal stress in lesions with foamy microglia compared to lesions with ramified microglia ^9^. This suggests that foamy microglia represents a state that contributes to demyelination, whereas ramified microglia supports endogenous myelin repair mechanisms. We compared the cellular and molecular profiles of mixed lesions containing ramified microglia versus lesions containing foamy microglia within the same MS donors. By this approach, we extracted the complex interplay between microglial in different states, their interactions within cellular micro-environment associated with lesion dynamics and outcome. The cellular resolution and completeness of our approach revealed that ramified microglia foster an environment conducive to repair and remyelination as they exhibited gene expression profiles indicative of myelin stability (*ABCA2*, *QKI*), while lesions with foamy microglia are associated with lesion expansion. Lesions with foamy microglia show increased immune activation, lymphocyte density, immunoglobulin (Ig) production (*IGHG1*, *IGHG3*), complement (*C1QB*, *C1QA*), iron dysregulation (*FTL*, *FTH1*), immune-oligodendrocytes, and demyelinating activity.

### Identification of the molecular signature of distinct cell niches

We performed high-resolution unbiased spatial transcriptomic profiling with large-field-of-view using Stereo-seq^6^ on post-mortem subcortical white matter (WM) mixed lesions with ramified microglia, mixed lesions with foamy microglia, and normal-appearing WM (NAWM) of MS donors (*n=8*) and subcortical WM of non-neurological control donors (*n=3*) (**Fig. 1A**) that were matched for age, sex, post-mortem delay, and pH of the CSF (**Table 1**).

**Figure 1:**
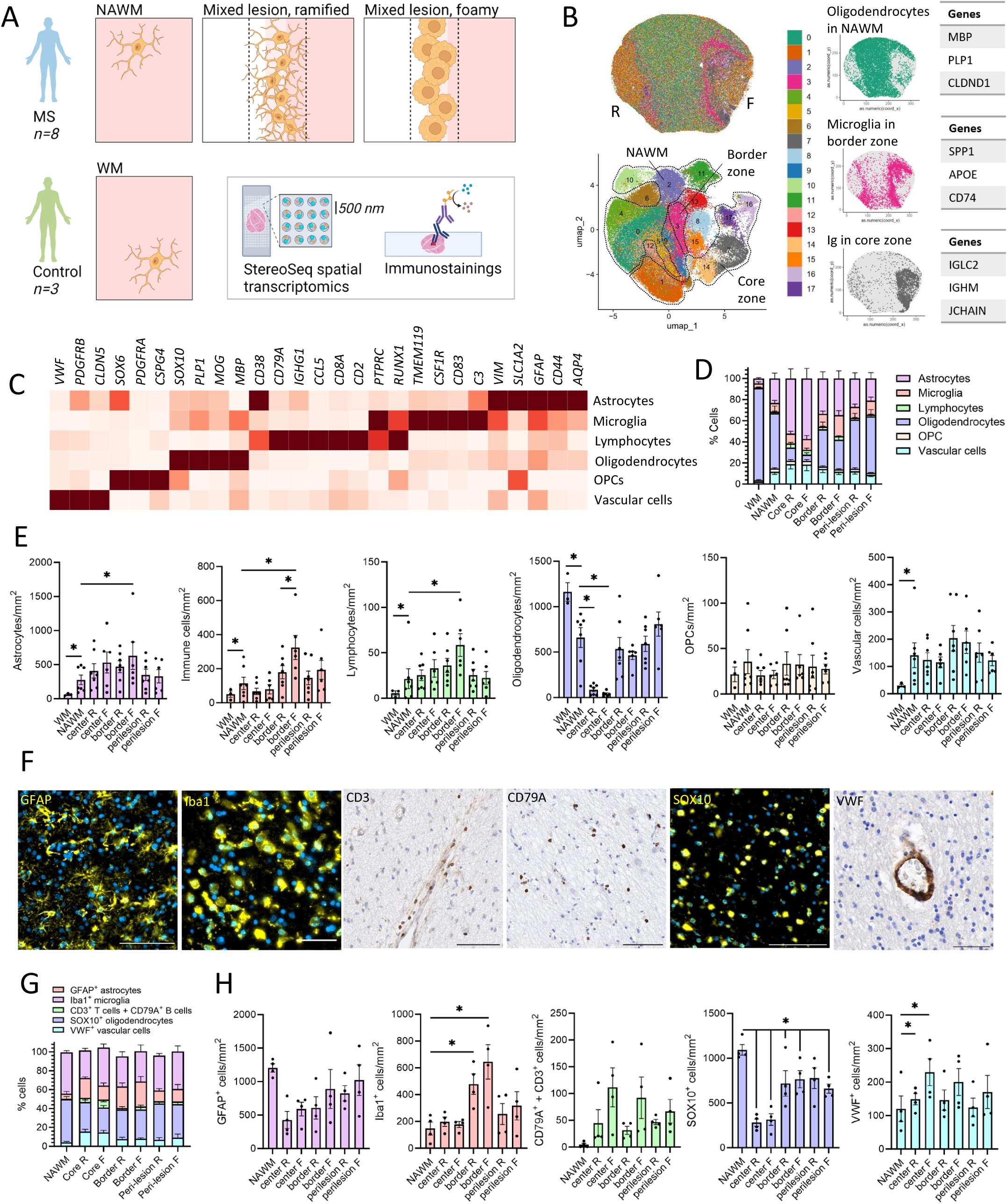
Cellular composition of mixed lesions with ramified and foamy microglia. A) Post-mortem human tissue of NAWM and mixed lesions with ramified or foamy microglia of MS donors and WM of control donors was collected for spatial transcriptomics and immunostainings. B) Representative image of one donor of unsupervised Seurat clustering of bins (25 µm) identified distinct clusters in NAWM, core, and border zones of mixed lesions, and distinguished between mixed lesions with ramified or foamy microglia. C) Cell bins were deconvoluted for major cell types astrocytes, microglia, lymphocytes, oligodendrocytes, OPCs, and vascular cells, and D) percentage and E) density of cell types was quantified. F) IHC of CD3 (scale bar 100 µm), CD79A (scale bar 100 µm), VWF (scale bar 50 µm), Iba1 (scale bar 50 µm), GFAP (scale bar 100 µm), and SOX10 (scale bar 100 µm), and their G) proportions and H) density. Significance was tested with a quasi-Poisson generalized linear model to compare the raw counts with offset for area or total number of cells, correcting for multiple testing with FDR. Significant differences indicated WM *vs* NAWM, for NAWM compared to all zones of interest, and for the cores, borders or peri-lesion zones of mixed lesions with ramified *vs* those with foamy microglia. *: p<0.10.

**Table 1:** Donor demographics of the spatial RNA sequencing cohort of MS and control samples.

| Donor | Group | MS type | Sex | Age (years) | PMD (hours) | pH CSF | Onset (years) | EDSS6 (years) | DOD (years) | Cause of death |
| --- | --- | --- | --- | --- | --- | --- | --- | --- | --- | --- |
| D5 | MS | SP | F | 43 | 10:45 | - | 22 | 16 | 21 | Subdural hematoma, pneumonia |
| D4 | MS | SP | F | 56 | 08:25 | 6.16 | 22 | 2 | 34 | Respiratory insufficiency by pneumonia |
| D6 | MS | PP | F | 54 | 09:20 | 6.27 | 23 | 8 | 31 | Heart failure |
| D3* | MS | SP | M | 50 | 10:50 | 6.55 | 29 | 17 | 21 | Euthanasia |
| D7 | MS | SP | F | 47 | 08:35 | 5.78 | 18 | 3 | 29 | Aspiration pneumonia |
| D1* | MS | PP | F | 67 | 11:25 | 6.80 | 38 | 7 | 29 | Pneumonia |
| D2* | MS | PP | M | 63 | 10:00 | 6.52 | 34 | 17 | 30 | Aspiration pneumonia, sepsis |
| D8* | MS | SP | F | 60 | 05:05 | 6.80 | 38 | 21 | 23 | Euthanasia |
| C1 | CON | NA | M | 79 | 06:20 | 6.65 | NA | NA | NA | Metastatic prostate cancer, dehydration |
| C2 | CON | NA | M | 57 | 04:55 | 6.51 | NA | NA | NA | Liver cirrhosis, urosepsis |
| C3 | CON | NA | F | 80 | 08:45 | 6.88 | NA | NA | NA | Electrolyte imbalance |

\*Donors with mirror formalin-fixed paraffin-embedded block available for validation. CON: control, DOD: duration of disease, EDSS6: years to EDSS6, F: female, M: male, NA: not applicable, Onset: age at onset, PMD: post-mortem delay, PP: primary progressive, SP: secondary progressive.

First, we applied a spot binning (bin50; 25-µm^2^) and clustering step to identify tissue domains representing local cell environments associated with pathological processes in MS tissue. With unsupervised Seurat clustering, we identified clusters associated with the NAWM, border zone, and core zone of mixed lesions. An overview of all clusters per donor and genes driving the clustering is provided in the **extended data file 1**. Clustering results of a representative donor are shown in **Fig. 1B**. Bin clusters that predominantly map to NAWM had high counts for oligodendrocyte and myelin-related genes (*MBP*, *PLP1*, *CLDND1*, *MOBP*, *MAG*, *APLP1*, *MOG*). Bins that map to mixed lesion cores were enriched for genes associated with an activated astrocyte phenotype, known as astrogliosis (*GFAP*, *S100B*, *VIM*, *AQP4*). In all donors, clusters associated with the core and the border zone of mixed lesions had enriched levels of Ig gene transcripts (*IGHGP*, *IGKC*, *IGHG3*, *IGHG1*, *IGHG4*, *IGHG2*, *IGLC3*, *IGLC2*, *IGKV3015*, *IGLC7*, *IGHM*, *IGLL5*, *JCHAIN*, *IGLV1-51*, *IGHGP*), which for some donors (3/7) was visually more evident in the core and border zone of mixed lesions with foamy microglia compared to lesions with ramified microglia. For a set of donors, bin clusters specific for the border zone of mixed lesion with foamy microglia could be identified, indicating the spatial and temporal complexity of MS lesion dynamics. These border zone of mixed lesion clusters had elevated levels of gene transcripts associated with microglia inflamed in MS (MIMS) or disease associated microglia (DAMS) ^10^ (*FTH1*, *HLA-B*, *APOE*, *SPP1*, *HLA-B*, *CD74*, *FTL*, *C1QB*, *HLA-DRA*, *C3*, *HLA-DRB1*, *GPNMB*, *C1QA*, *TREM2*). In addition, the border zone of mixed lesions with foamy microglia were also often enriched with gene transcripts associated with astrogliosis. Donors with clusters associated with Ig-genes in NAWM also had clusters associated with MIMS or DAMS and astrogliosis in NAWM. For another set of donors, bin clusters associated with MIMS or DAMS were more sparsely distributed throughout the border zone, peri-lesional zone, and NAWM. This approach successfully segmented tissue domains and zones with distinct cellular and molecular features. The molecular signatures of these zones confirm many of the reported molecular and cellular changes associated with myelination state, microglia states, and Ig production in mixed lesions ^2,11,12^, and additionally highlights differences between lesions with different microglial states. The high resolution and preservation of spatial information of our assays revealed donor and lesion state heterogeneity.

### Cell type composition of mixed lesion zones containing ramified or foamy microglia

Cell type composition of mixed lesions was previously characterized ^10^. However it is unclear whether cell type composition varies for lesions with different microglial states. Therefore, we determined the cell type composition of cell bins based on nuclei staining within tissue zones of NAWM and mixed lesions with ramified or foamy microglia in MS tissue and of WM in control tissue. First, manual segmentation of tissue zones was performed based on expression of HLA-related genes, density and morphology of HLA^+^ cells, expression of *MBP* and *PLP*, and RNA density. Next, single cell segmentation was performed using the CellBin pipeline ^13^, and major cell types (astrocytes, microglia cells, lymphocytes, oligodendrocytes, oligodendrocyte precursor cells (OPCs) and vascular cells) were annotated using RCTD and the single-nucleus RNA-sequencing dataset of Absinta *et al.* ^10^ (**Fig. 1C-E**). Results were validated with immunohistochemistry (IHC) of CD3 for T cells, CD79A for B cells, VWF for endothelial cells, Iba1 for microglia/macrophages, GFAP for astrocytes, and SOX10 for oligodendrocytes (**Fig. 1F-H**). For all false-discovery rate (FDR) corrected p-values see **extended data file 2**.

Compared to control WM, MS NAWM exhibits a higher density of astrocytes, immune cells, lymphocytes, and vascular cells, and a relatively lower density of oligodendrocytes. The density of OPCs was comparable between WM and NAWM. Within MS donors, the density of astrocytes, immune cells, lymphocytes, OPCs, and vascular cells was comparable for peri-lesion zones and NAWM. In the peri-lesion region of mixed lesions with foamy microglia compared to the NAWM, the density of oligodendrocytes was lower (only when assessed with IHC). This was not found for the peri-lesion zone of mixed lesions with ramified microglia. In the border zones of mixed lesions, the density of OPCs and vascular cells was comparable for border zones and NAWM. The density of immune cells was higher compared to the NAWM, and a higher density of immune cells was found in mixed lesions with foamy microglia compared to those with ramified microglia. Specifically in the border of mixed lesions with foamy microglia compared to the NAWM, a higher density of astrocytes and lymphocytes was found, and a lower density of oligodendrocytes. In the core zones of mixed lesions, compared to the NAWM, a relative increase of astrocytes, a higher density of vascular cells, and a lower density of oligodendrocytes were found. Specifically in the core zone of mixed lesions with foamy microglia compared to NAWM, a higher density of lymphocytes was found. This cellular profile aligns with previously reported characteristics of NAWM and mixed lesions ^10^, which validates the segmentation and deconvolution method. The preservation of OPCs suggests that a differentiation block contributes to remyelination failure in MS. In sum, mixed lesions with foamy microglia are identified by a larger accumulation of microglia and larger infiltration of lymphocytes compared to mixed lesions with ramified microglia.

### Perivascular lymphocytes are more reactive compared to parenchymal lymphocytes

In MS, reactivated B and T cells accumulate in the CNS, and low numbers infiltrate the parenchyma from the perivascular space. These lymphocytes are involved in key pathogenic events that contribute to the pathogenesis of MS ^14^. Previously it is reported that lower proportion of parenchymal T cells contain TNF compared to perivascular T cells based on immunohistochemistry ^15^. This implies that perivascular lymphocytes are more reactive than parenchymal lymphocytes ^16^. Therefore, we set out to test this hypothesis by investigating the gene expression profile of parenchymal and perivascular lymphocytes using pre-ranked gene-set enrichment analysis (GSEA). On average, 211 ± 152 T cells and 129 ± 93 B cells were identified per sample. We annotated T and B cells that were in close proximity (<50 µm) to a cell annotated by the vascular cluster or clustered together (>3) as perivascular cuffs, and considered them perivascular lymphocytes (77.1% and 79.9%, respectively). The other lymphocytes were considered parenchymal T and B cells (22.9% and 20.1%, respectively) (**Fig. 2A**). The distribution of perivascular and parenchymal lymphocytes is in line with previous reported analysis of MS tissue ^17^. The number of perivascular lymphocytes was higher than the number of parenchymal lymphocytes in NAWM, border zones, core zones, and peri-lesional zones. In the border zone of mixed lesions with foamy microglia but not in the border zone of those with ramified microglia, perivascular lymphocyte cell density was increased compared to NAWM **(Fig. 2B)**. Gene ontology (GO) analysis was performed to predict functional changes based on molecular profiles. All enriched GO-terms are provided in **extended data file 3**, and a selection of most significant and MS relevant biological processes is visualized in **Fig. 2C**. Compared to perivascular lymphocytes, parenchymal lymphocytes had higher gene counts for functions associated with opsonization. On the contrary, compared to parenchymal lymphocytes, perivascular lymphocytes had higher gene counts for functions associated with T-cell activation and with generation and maintenance of tissue-resident memory T cells and T-helper cells. Additionally, compared to parenchymal lymphocytes, perivascular lymphocytes were enriched for functions indicative of B-cell activation, Ig production, and somatic diversification and recombination of Igs. Production of soluble mediators by activated perivascular lymphocytes has been postulated to contribute to lesion expansion ^16^. These results for the first time demonstrate the molecular transformation of lymphocytes while migrating into the parenchyma.

**Figure 2:**
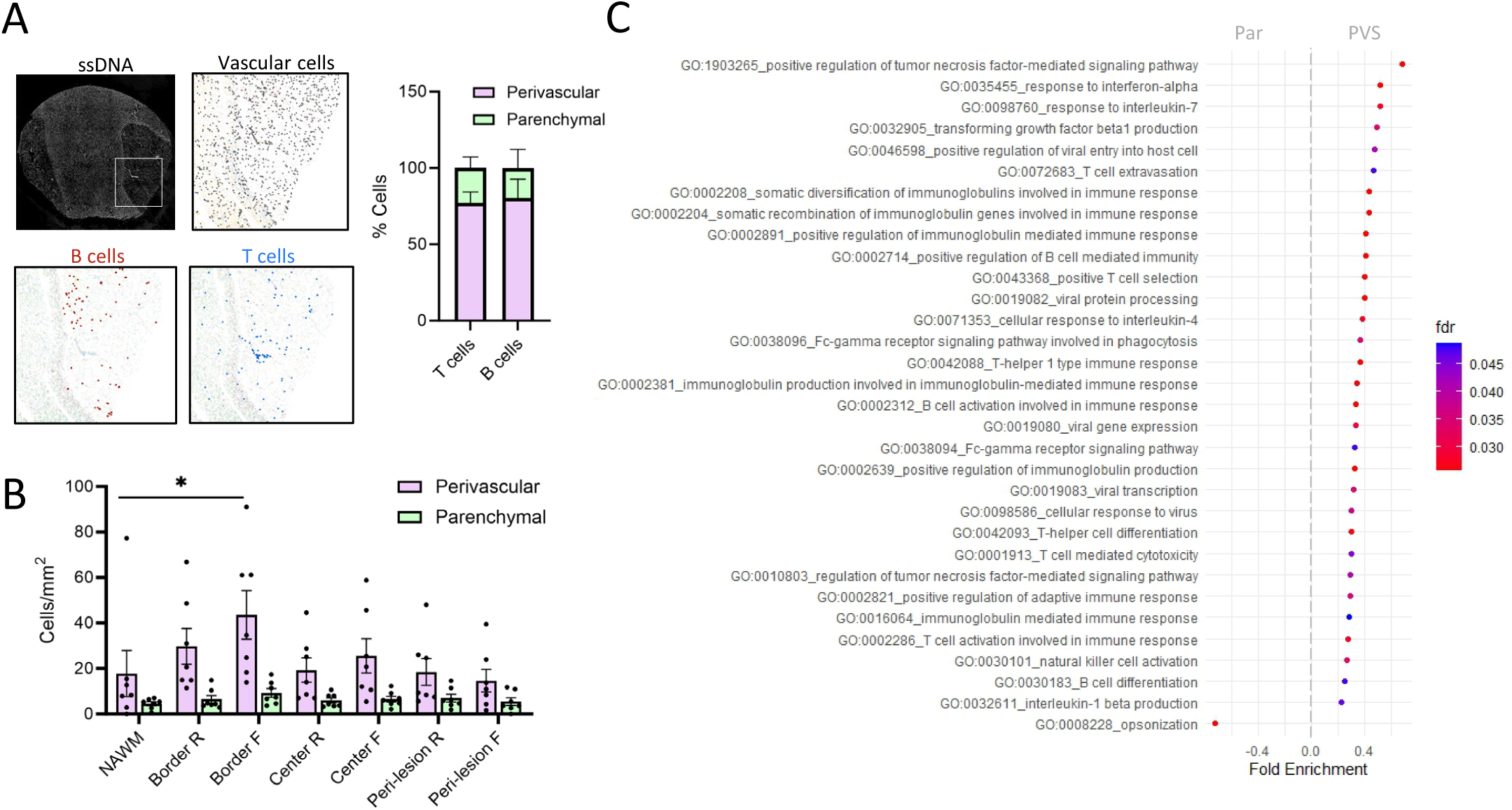
Perivascular lymphocytes are more reactive than parenchymal lymphocytes. A) Cell bins identified as vascular cells, T cells or B cells were used to quantify the number of perivascular and parenchymal lymphocytes. The majority of T and B cells were perivascular, as they were in close proximity (<50 µm) to a cell annotated by the vascular cluster or clustered together (>3) as perivascular cuffs. B) There was an enrichment of perivascular lymphocytes in the border of zone of lesions with foamy microglia. C) Pre-ranked GSEA of perivascular (PVS) lymphocytes compared to parenchymal (Par) lymphocytes.

### Cell states at the border of mixed lesions with ramified or foamy microglia

Microglial state can imply functional implications for lesion dynamics. We investigated microglial cell states associated with pathological zones of mixed lesions with ramified or foamy microglia. The immune cluster was further deconvoluted based on the single-nuclei RNA sequencing dataset of Absinta *et al*. (2021) ^10^. The relative population density of the microglial cells in WM, NAWM, and pathological zones is shown in **Fig. 3A**. Compared to control WM, the NAWM is enriched with macrophages and of MIMS with high expression of iron-associated genes (MIMS-iron). Previously, the density of MIMS was found upregulated in the border zone of mixed lesions ^10^. In the border zone of mixed lesions with foamy microglia compared to the NAWM, the density of MIMS with upregulation of genes associated with lipid scavenging and metabolism (MIMS-foamy) and of MIMS-iron was higher, which was not found in the border zone of mixed lesions with ramified microglia. Accordingly, the density of MIMS-iron was higher in the border zone of mixed lesions with foamy microglia compared to the border zone of mixed lesions with ramified microglia. Within each donor, the density of MIMS-iron is higher in the border zone of the lesion with foamy microglia compared to lesions with ramified microglia, as shown in **Fig. 3B** where the MIMS-iron is projected onto the cell bins, highlighting the robustness of this finding. These findings were confirmed using IHC, revealing a higher density of MIMS-iron in the border zone of mixed lesions with foamy microglia compared to the border zone of those lesions with ramified microglia was validated (**Fig. 3C**). Iron rim-positive lesions detected by MRI are associated with a higher clinical severity ^18^. Our findings indicate that mixed lesions with foamy microglia have more iron accumulation than mixed lesions with ramified microglia and may therefore be more aggressive.

**Figure 3:**
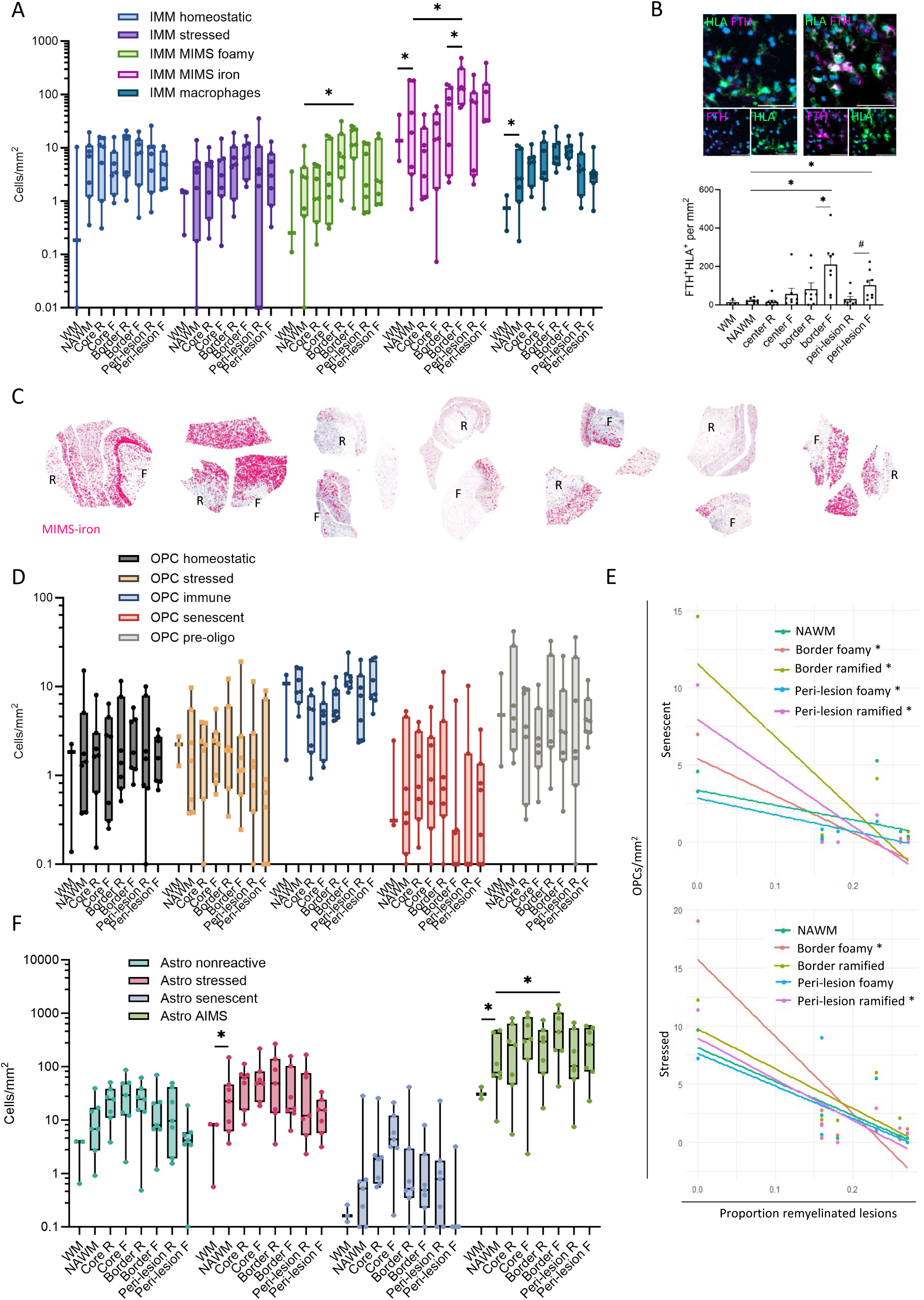
Cell states are associated with lesion expansion and failed remyelination. Further deconvolution of A) the immune cell cluster into cell states. B) IHC of HLA and FTH of a lesion with ramified microglia (left) and a lesion with foamy microglia (right) (scale bar 50 µm). There are more HLA^+^FTH^+^ microglia/mm^2^ in the border zone of mixed lesions with foamy microglia than in the border zone of mixed lesions with ramified microglia. C) MIMS-iron projected onto cell bins for all donors, lesions with foamy microglia indicated with F, those with ramified microglia indicated with R. D) Further deconvolution of the OPC cell cluster into cell states. E) Senescent and stressed OPC density negatively correlated to the proportion of remyelinated lesions. F) Further deconvolution of the astrocyte cell cluster. Significance of count data was tested with a quasi-Poisson generalized linear model to compare the raw counts with offset for area or total number of cells, correcting for multiple testing with FDR. Significant differences indicated for NAWM compared to all zones of interest, and for the cores, borders or peri-lesion zones of mixed lesions with ramified *vs* those with foamy microglia. Correlations were tested with Pearson’s correlation test, corrected with FDR. *: p<0.10.

As the relative density of OPCs is unaltered across the pathological zones in MS tissue, impaired OPC differentiation into myelinating oligodendrocytes could contribute to remyelination failure ^19^. Therefore, we further deconvoluted the OPC cell cluster into homeostatic, stressed, immune-like, senescent, and pre-oligo OPCs. The projected density of OPC cell states on the MS tissue was then correlated to proportion of remyelinated lesions within each donor. The densities of OPC cell states in the WM, NAWM, cores, border zones, and peri-lesional zones of control tissue and mixed lesions with ramified or foamy microglia were comparable (**Fig. 3D**). The density of senescent and stressed OPCs, however, negatively correlated to the overall proportion of remyelinating lesions identified at autopsy. (**Fig. 3E**, p-values shown in **extended data file 4**). Our data indicate that donor-specific impaired OPC differentiation and maturation caused by cellular senescence and oxidative stress ^20,21^ may contribute to impaired remyelination and thus underly the high variation of remyelinated lesions among people with MS.

Previously, a subcluster of astrocytes, the astrocytes inflamed in MS (AIMS), was identified in the border of mixed lesions that exhibit neurodegeneration associated profiles and cross-talk with MIMS ^10^. To investigate the density of AIMS in pathological zones of mixed lesions with ramified or foamy microglia, we further deconvoluted the astrocyte cluster into cell-states. The density of astrocyte cell-states are visualized in **Fig. 3F**. In NAWM, we identified an enrichment of reactive astrocytes and AIMS compared to control WM. Within MS donors, compared to NAWM an enrichment of AIMS was identified in the border of mixed lesions with foamy microglia but not in the border of those with ramified microglia. Together, the higher density of lymphocytes, MIMS, and AIMS in mixed lesions with foamy microglia compared to those with ramified microglia, indicates that those with foamy microglia are more degenerative and inflammatory, and therefore more aggressively inducing cell state shifts in astrocytes.

### Receptor–ligand pair interactions associated with lesion expansion

Utilizing the preserved spatial location of single cells, we investigated the interactome of cell states in pathological zones. Communicating cells were identified through upregulation of ligand–receptor pairs of in adjacent cells (<100 μm) and nearby cells (<200 μm) using Cell-Chat. Ligand–receptor pairs per donor for each zone is provided in **extended data file 5**. The top number of interactions as well as the highest strength of interactions were found in the border zone of lesions with foamy microglia. Stressed oligodendrocytes were the main signal providers, and stressed oligodendrocytes, MIMS-iron, and immune-OPCs were the main signal receivers (**extended Fig. 1**). Receptor–ligand pairs that functionally may indicate lesion expansion are summarized in **Table 2**. Increased receptor–ligand interactions indicative of oxidative damage, demyelination, and lymphocyte activation were identified in the border zone of mixed lesions with foamy microglia compared to the border zone of those with ramified microglia. Less receptor–ligand interactions indicative of T-cell inhibition were identified in the peri-lesional zone of mixed lesions with foamy microglia compared to the peri-lesional zone of those with ramified microglia. Together, this further highlights the destructive interplay of lymphocytes activation and oxidative stress driving pathological demyelination and indicates that lesions with foamy microglia are more prone to lesion expansion than lesions with ramified microglia.

**Table 2:** Number of receptor-ligand pair interactions in the border and peri-lesion zone of mixed lesions with ramified or foamy microglia.

| Receptor–ligand pair | Interpretation | Donors | Border |  | Peri-lesion |  |
| --- | --- | --- | --- | --- | --- | --- |
|  |  |  | R | F | R | F |
| APP–CD47 | Failed cell repair after oxidative damage<sup>22</sup> | 7/7 | 98 | 204 | 91 | 98 |
| PSAP–GPR37 | Lysosomal degradation of sphingolipids<sup>23</sup> | 5/7 | 52 | 127 | 42 | 52 |
| Cholesterol–<br>cholesterol–LIPA–<br>RORA | Inflammatory processes and differentiation of Th17 cells<sup>24</sup> | 4/7 | 19 | 62 | 19 | 21 |
| APOE–TREM2–<br>TYROBP | Lipid metabolism and phagocytosis<sup>25</sup> | 2/7 | 22 | 56 | 22 | 20 |
| PSAP–GPR37L1 | Lipid metabolism and phagocytosis<sup>23</sup> | 5/7 | 10 | 76 | 10 | 19 |
| APP–TREM2–TYROBP | Lipid metabolism and phagocytosis<sup>25</sup> | 2/7 | 18 | 43 | 18 | 26 |
| PTN–NCL | Lymphocyte survival and recruitment<sup>26,27</sup> | 3/7 | 2 | 31 | 2 | 9 |
| CD99–CD99 | Lymphocyte survival and recruitment<sup>26,27</sup> | 5/7 | 4 | 16 | 4 | 9 |
| SEMA4D–PLXNB1 | Repulsion of axonal growth cones<sup>28</sup> | 3/7 | 2 | 6 | 2 | 4 |
| C3–ITGAX–ITGB2 | Complement mediated activation of microglia<sup>29</sup> | 1/7 | 0 | 0 | 0 | 10 |
| SPP1–CD44 | Inhibition of T-cell activity<sup>30</sup> | 5/7 | 117 | 129 | 117 | 65 |

R: ramified, F: foamy

### Gene profile of lesion expansion and remyelination

Our study demonstrates that mixed lesions with foamy microglia are more prone to lesion expansion, while those with ramified microglia are more likely to transit into inactive or remyelinated lesions. Direct comparisons of RNA expression in pathological zones between these distinct microglial states enabled us to identify gene expression profiles linked to lesion expansion and repair.

We conducted single-cell level analyses comparing control WM with NAWM as well as the border and peri-lesion zones of mixed lesions with foamy microglia versus ramified microglia. **Fig. 4A-C** shows volcano plots visualizing differentially expressed (DE) genes, and enriched gene sets potentially related to lesion expansion, repair, and remyelination are highlighted in **Fig. 4D-E**. The top-200 DE genes are listed in **extended data file 6**, and all gene sets enriched in the border or peri-lesion zones are listed in **extended data file 7**. To understand the involvement of specific cell types in lesion dynamics, we mapped DE genes onto the UMAP of the single-cell dataset from Absinta *et al*. ^10^ **(extended Fig. 2)**.

**Figure 4:**
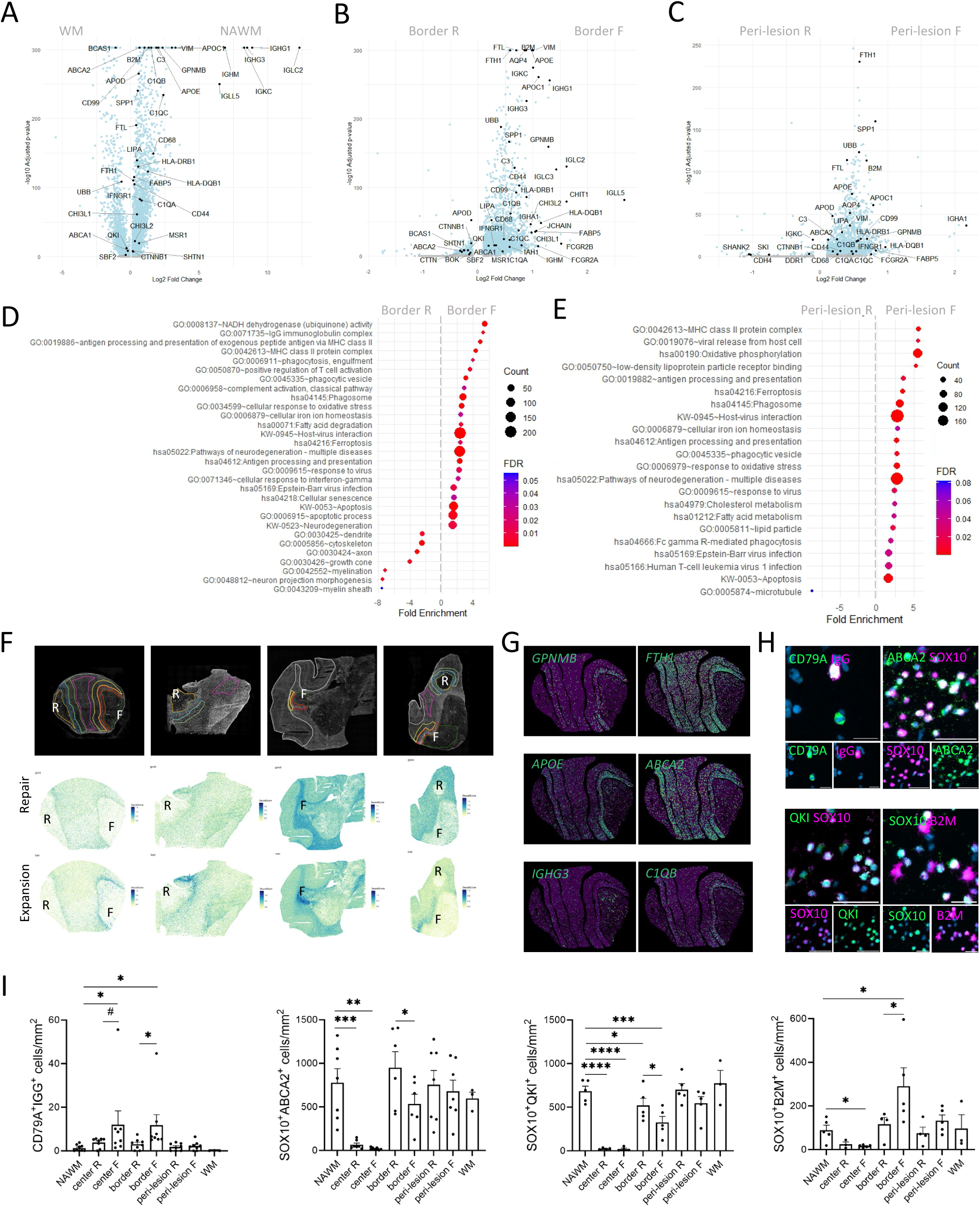
Gene expression patterns associated with lesion expansion and repair and immunohistochemical validation. Volcano plots of A) WM compared to NAWM, B) border zones of mixed lesions with foamy microglia compared to those with ramified microglia, C) peri-lesion zone of mixed lesions with foamy microglia compared to those with ramified microglia. Highlighted gene sets likely associated with lesion expansion or repair in D) the border zone of mixed lesions with foamy microglia compared to those with ramified microglia, and E) in the peri-lesion zone of mixed lesions with foamy microglia compared to those with ramified microglia. F) Gene modules associated with lesion expansion or lesion repair are projected on cell bins. G) Genes of interest projected on cell bins. H) IHC double stainings of CD79A + IgG (scale bar 25 µm), SOX10 + ABCA2 (scale bar 50 µm), SOX10 + QKI (scale bar 50 µm), and SOX10 + B2M (scale bar 50 µm). I) There are more CD79A^+^IgG^+^ cells/mm^2^ in the border zone and core of mixed lesions with foamy microglia than in the border zone and core of mixed lesions with ramified microglia, less SOX10^+^ABCA2^+^ oligodendrocytes/mm^2^ and less SOX10^+^QKI^+^ oligodendrocytes/mm^2^ in the border zone of mixed lesions with foamy microglia than in the border zone of mixed lesions with ramified microglia, and more SOX10^+^B2M^+^ oligodendrocytes/mm^2^ in the border zone of mixed lesions with foamy microglia compared to the border zone of mixed lesions with ramified microglia. Significance was tested with a quasi-Poisson generalized linear model to compare the raw counts with offset, correcting for multiple testing with FDR. #: p<0.11, *: p<0.10, **: p<0.01, ***: p<0.001, **** p<0.0001.

In NAWM, 2,386 genes showed increased counts compared to healthy WM, particularly those associated with oligodendrocytes, astrocytes, and immune cells. In line with previous reports ^31,32^, gene counts were indicative of activation of the complement cascade, microglia activation, cytokine production, lipid metabolism, phagocytosis, Ig production, and iron dysregulation (**Fig. 4A**). This suggests that microglial and astrocyte activation, along with oligodendrocyte vulnerability, precede lesion formation. In border zones of lesions with foamy microglia, 2,518 genes had higher counts and 535 had lower counts compared to those with ramified microglia. Genes with lower counts in lesions with foamy microglia were associated with oligodendrocytes and OPCs, suggesting greater oligodendrocyte destruction and reduced remyelination capacity. In peri-lesion zones, lesions with foamy microglia showed higher counts for 1,768 genes, which associated with oligodendrocytes, astrocytes, and microglia, suggesting greater oligodendrocyte vulnerability and astrocyte/microglial activation, which could contribute to lesion expansion. Many DE genes were shared between border and peri-lesional zones of lesions with foamy microglia (1,377 genes), with fewer genes shared in lesions with ramified microglia (22 genes). Overlapping genes in border and peri-lesional zones were associated with phagocytosis, lipid metabolism, immune activation, immune-oligodendrocytes, iron dysregulation, and neurodegeneration, supporting their role in lesion expansion ^10,33^. Conversely, gene sets related to axonal growth and myelination were enriched in border zones of lesions with ramified microglia, indicating a higher degree of repair and possibly remyelination **(Fig. 4B-E)**.

Gene modules related to remyelination and axonal outgrowth (*APOD*, *CTNNB1*, *QKI*, *ABCA2*, *SBF2*, *BOK*, *CTTN*, *SHTN1*, *BCAS1*, *SHANK2*, *CDH4*, *SKI*, *NEAT1*, *DDR1*) and related to lesion expansion (*APOC1*, *APOE*, *APP*, *B2M*, *C1QA*, *C1QB*, *C1QC*, *C3*, *CD44*, *CD68*, *CD74*, *CD99*, *CHI3L1*, *CHIRL2*, *CHIT1*, *CLU*, *FABP5*, *FCGR1A*, *FCGR2A*, *FCGR2B*, *FTH1*, *FTL*, *GPNMB*, *HLA-A*, *HLA-DQB1*, *HLA-DRB1*, *IAH1*, *IFNGR1*, *IHA1*, *IGHG1*, *IGHG3*, *IGHM*, *IGLC2*, *IGLC3*, *IGLL5*, *JCHAIN*, *LIPA*, *MSR1*, *NCAM1*, *OLR1*, *DERPINA3*, *SPP1*, *UBB*) were generated and projected onto the cell bins (**Fig. 4F**, for all donors see **extended Fig. 3**). The gene module related to remyelination and axonal outgrowth was expressed at lower levels in lesions with foamy microglia **(Fig. 4F-G)**. Accordingly, our IHC analysis validated this at the protein level, showing a loss of QKI^+^SOX10^+^ oligodendrocytes, necessary for mature myelin maintenance ^34^, in the border zone of mixed lesions with foamy microglia compared to those with ramified microglia, and lower density of ABCA2^+^SOX10^+^ oligodendrocytes in the border zone of mixed lesions with foamy microglia compared to those with ramified microglia. This indicates dysregulated sphingolipid metabolism ^35^ **(Fig. 4H-I)**. Together, this reflects decreased axonal receptiveness to remyelination ^36^ in the border zone of mixed lesions with foamy microglia, and highlights the role of oligodendrocytes in perpetuated demyelinating activity and loss of myelin stability and integrity.

In contrast, the gene module linked to lesion expansion was more abundant in lesions with foamy microglia **(Fig. 4F-G)**. We validated our findings at protein level, through IHC of IgG and B2M **(Fig. 4H-I)**. The border zone of lesions with foamy microglia was identified by a higher density of B2M^+^SOX10^+^ oligodendrocytes compared to NAWM and compared to the border zone of lesions with ramified microglia **(Fig. 4H-I)**, indicating a higher number of immune-oligodendrocytes. In neurodegenerative diseases, OPCs and oligodendrocytes can present MHC class I and II antigens, which can be induced by IFNƴ. In MS, these immune OPCs and immune oligodendrocytes act as active immunomodulators and cytotoxic targets, thereby contributing to a perpetuated autoimmune response and ongoing oligodendrocyte destruction and consequently myelin degradation ^37,38^. Furthermore, the number of IgG^+^CD79A^+^ B cells was higher in the border zone of mixed lesions with foamy microglia compared to the border zone of those with ramified microglia, and there was a trend in the core. The increased level of IgG production by CD79A^+^ B cells in the border zone and core of mixed lesions with foamy microglia compared to those with ramified microglia may play a role in sustained neuro-inflammation (**Fig. 4H-I**). These finding expand earlier work showing IgG-related gene transcription in the rim of mixed lesions ^12^ and the association of lesional and meningeal CD20^+^ B cells with CSF oligoclonal IgG production ^39^. Igs can break microglial immune-tolerance ^40^ and activate the complement cascade ^41^, contributing to sustained inflammation and lesion expansion. Our findings indicate upregulation of complement-related genes and complement-mediated activity in the border zone and peri-lesional zone of mixed lesions with foamy microglia, suggesting a role of immune-complex formation and related complement activation in lesion expansion.

## Discussion

Microglia are highly dynamic cells integral to critical CNS functions, including myelinogenesis, phagocytic debris clearing, myelin and synaptic remodeling, and immune surveillance. These cells undergo intricate morphological, transcriptional, and functional changes in response to environmental stimuli and due to phagocytosis ^7^. In MS, microglia play a crucial role in lesion dynamics. Mixed lesions can expand, become inactive, or undergo remyelination, however the molecular and cellular mechanisms underlying these processes remain poorly understood. In this study, we generated capture based high-resolution spatial transcriptomics profiles of post-mortem human MS tissue, to investigate the cellular composition, gene expression profiles, and intercellular communication in mixed lesions with ramified microglia and of those with foamy microglia within the same MS donors and compared to the NAWM of control donors. Our findings indicate that mixed lesions with ramified microglia are more likely to transit into inactive or remyelinated lesions, whereas mixed lesions with foamy microglia tend to exhibit continuous lesion activity and undergo degeneration, highlighting how microglia can mediate lesion dynamics. In addition we were able to identify molecular and cellular features associated with lesion fate.

In mixed lesions with ramified microglia, pathways involved in axonal protection and myelin stability combined with lower complement and inflammatory responses likely safeguard surrounding tissue from lesion expansion. Notably, in the rat brain, QKI identifies actively myelinating oligodendrocytes ^42^. The preservation of QKI^+^ oligodendrocytes in mixed lesions with ramified microglia suggests enhanced remyelinating potential. Additionally, ABCA2 is involved in low-density lipoprotein trafficking and is crucial for regulation of the sphingolipid metabolism and therefore myelin integrity ^35,43^. Maintained population of ABCA2^+^ oligodendrocytes in mixed lesions with ramified microglia therefore contributes to myelin stability and resistance to ongoing demyelination. In mixed lesions, activated B and T cells likely contribute to the cell state change of microglia from ramified to foamy, by producing Igs and pro-inflammatory cytokines. Igs can opsonize myelin ^11^, activate the complement cascade ^41^, and disrupt microglial immune-tolerance ^40^, which sustains microglial activation and demyelination. Pro-inflammatory cytokines further induce immune-oligodendrocytes, which become cytotoxic targets ^37,38^, leading to their destruction and subsequent iron release. Microglia clear up this iron, but its uptake inhibits lysosomal acidification ^44^, impacting on the metabolism of phagocytosed myelin. Iron-laden microglia produce high levels of reactive oxygen species (ROS), causing oxidative damage to DNA, lipids and proteins ^31^. Oxidized lipids perpetuate inflammation ^45^. Additionally, ROS-induced oxidative stress and inflammation can induce stressed or senescent OPCs that have a differentiation block ^46^, leading to failed remyelination. Iron chelators, such as deferoxamine, might serve as promising therapeutic targets to suppress iron-mediated lesion expansion of lesions with foamy microglia.

These zone- and donor-specific mechanisms of degeneration and repair failure offer insights in mechanisms underlying lesion dynamics. We furthermore highlight the importance of microglial phenotyping for future studies on MS lesions. Understanding the distinct roles of microglial states in lesion dynamics is crucial for developing targeted interventions to modify the disease course in MS.

## Acknowledgements

We are grateful to the brain donors and their families for their commitment to the Netherlands Brain Bank donor program. We acknowledge the DCS Cloud platform (https://eu-cloud.stomics.tech) to provide workflows and tools for omics data analysis, interactive visualization, and collaborative project and data management.

## Methods

### Donors

Post-mortem subcortical WM tissue of n = 8 MS donors (6 females, 2 males) and n = 3 healthy control donors (1 female, 2 males) was provided by the Netherlands Brain Bank (Amsterdam, The Netherlands, www.brainbank.nl). All donors provided informed consent for brain autopsy and the use of their tissue and clinical data for research purposes in compliance with national ethics guidelines. For all MS donors, MS pathology was confirmed by a certified neuropathologist. Donor demographics are shown in **Table 2**. The cohort was balanced for sex, age, post-mortem delay, and the pH of the CSF. For control donors, fresh-frozen subcortical WM was collected (n = 3). For MS donors, fresh-frozen subcortical WM lesions were collected (n = 8). If available, formalin-fixed paraffin-embedded (FFPE) mirror blocks were additionally collected (n = 4). MS lesions were characterized as previously described ^3,5,47^. Mixed lesions had a hypo-cellular and fully demyelinated core. At the border zone of the lesion, there was an accumulation of HLA-DR^+^ microglia. For each donor, one mixed lesion with ramified microglia and one mixed lesion with foamy microglia was included.

### Tissue preparation and spatial sequencing

Stereo-seq samples were prepared by flash-freezing post-mortem human brain tissue at -80°C. Tissue was processed for Stereo-seq as previously described ^6^. In brief, 10-μm cryosections were collected and adhered to Stereo-seq chips (up to 2cm x 3cm) and fixed in methanol at -20°C for 30 minutes. The chips were permeabilized at 37°C for 12 to 18 minutes and washed with 0.1 × SSC buffer containing 0.05 U/μl RNase inhibitor. RNA captured by the DNA nanoball on the chips was reverse transcribed at 42°C for 180 min. cDNA was then purified and amplified using PCR. Stereo-seq libraries were prepared as previously described by Chen et al. (2022) ^6^ and sequenced on an MGI DNBSEQ-Tx sequencer.

### Stereo-seq data processing

Raw sequencing data were processed using the Stereo-seq Analysis Workflow (SAW) v5.1.4 ^48^ integrated on the DCS Cloud platform (https://eu-cloud.stomics.tech). Briefly, CID sequences from read 1 were mapped to the Chip T mask file allowing 1 mismatch. Unqualified MID reads were discarded, and short reads (less than 30 bases after trimming) filtered. The resulting clean reads were mapped to the hg38 reference genome to generate a gene-spot expression matrix, for which counts can be aggregated to different square-bin sizes using a customized script in R.

Raw counts at bin 50 level were processed using the *sctransform* function in Seurat v5.0.1 ^49^ by regressing out mitochondrial expression, *RunUMAP* with dims=1:20 and *FindClusters* with a resolution of 0.5. This was performed for each individual Stereo-seq chip including all bin units. Cluster markers were calculated using the FindAllMarkers function in Seurat with default method and used to check differential expression between clusters.

### Cell segmentation and cell type annotation

Cell segmentation was performed based on the single-strand DNA image using the CellBin pipeline ^13^. The GMM method was applied for cell correction afterwards. Segmented cells were filtered to have minimal gene and MID counts of 10 and mitochondrial expression <20%. For cell type annotation, we only kept cells in the outlined zones (see below methods) and used the single-cell RNA-sequencing data from Absinta *et al*. ^10^ as a reference for cell type annotation. Briefly, we removed chronic inactive and neuron data from the scRNA-seq dataset and constructed a reference using the Reference function in spacexr v2.2.1 (previously named RCTD ^50^). Major cell types (astrocytes, microglia, lymphocytes, oligodendrocytes, oligodendrocyte precursor cells, and vascular cells) were deconvoluted to the Stereo-seq spatial data using the run.RCTD command in “doublet” mode. The majority of cells in the immune cluster of Absinta *et al*. ^10^ are microglia, therefore the immune cluster was renamed as microglia cluster. The cell type of a cell bin spot was assigned with the largest weight. For annotation of subtypes, we subdivided the single-cell and spatial datasets and ran RCTD separately for each cell type using the same settings. Cell density in each zone in the Stereo-seq data was calculated using cell number and zone area, which was calculated roughly as number of bin50 units x 25 x 25 μm^2^.

### Differential gene expression in zones of interest

To calculate differential gene expression between outlined zones, we merged the data from all donors and used Seurat to normalize the counts and ran FindMarkers with default parameter. Genes adjusted p-value <0.05 were regarded as differentially expressed genes (DEGs).

### Characterization of perivascular and parenchymal lymphocytes

A parenchymal lymphocyte is characterized by the lack of vascular cells and less than 3 neighboring lymphocytes in a 50-μm radius. The adjacency matrix within 50-μm radius was calculated using the getAdj_manual function from the DR.SC package (v3.3 ^51^). To calculate differential expression between perivascular and parenchymal types, we used the FindMarker function from Seurat using the default Wilcoxon method. We also aggregated the counts per type per donor and fed the pseudo-bulk counts to DESeq2 v1.40.2 ^52^ and calculated corresponding log2FoldChange and padj values, which were used for gene set enrichment analysis (GSEA) using the gseGO function from ClusterProfiler v4.8.3 ^53^. We set the ontology to “BP”, minGSSize to 10, and maxGSSize to 500.

### Cell–cell interaction using CellChat

Cell–cell interactions within a specific zone were predicted using CellChat v2.1.2 ^54^ at subtype levels. We used the integrated CellChatDB for ligand-receptor interactions in human and performed computeCommuProb using the following parameters: *trim = 0.1, distance.use = TRUE, interaction.range = 250, scale.distance = 0.1, contact.dependent = TRUE, contact.range = 100.* Communication pathways were kept for those with minimal 10 cells. Communication networks were visualized using circle plots or heatmaps within CellChat. Ligand–receptor pairs contributing to the pathways were retrieved using the netAnalysis_contribution command.

### Immunohistochemistry

For immunohistochemistry, 8-µm sections of FFPE tissue were deparaffinized and rehydrated in a xylene and ethanol series. Antigen retrieval was performed using citrate buffer or EDTA buffer (**extended data file 8**). Fresh-frozen sections were cryo-sectioned at 10 µm and fixated. Sections were incubated with 1% H_2_O_2_ in phosphate buffered saline (PBS) + 0.5% TritonX for 20 minutes. Non-specific antibody binding was blocked with blocking buffer (PBS + 10% normal horse serum + 1% bovine serum albumin + 0.5% TritonX) for 1 hour. Primary antibodies were incubated overnight at 4°C, at dilutions indicated in **extended data file 8**. For 3,3’-diaminobenzidine (DAB) stainings, appropriate secondary antibodies 1:400 in blocking buffer were incubated for 1 hour, and avidin-biotin complex 1:800 in PBS was incubated for 45 minutes. DAB envision kit 1:100 was incubated to visualize the staining and sections were counterstained with hematoxylin before dehydration and coverslipped with Entellan. For immunofluorescent stainings, sections were either incubated with the appropriate secondary antibodies conjugated to fluorophores 1:800 for 1 hour, or incubated with appropriate biotinylated secondary antibodies 1:400, avidin-biotin complex 1:800 for 45 minutes, biotinylated tyramide 1:10,000 for 10 minutes and streptavidin conjugated fluorophore 1:800 for 1 hour. Nuclei were stained with DAPI 1:1,000 before incubation with Sudan black 0.1% for 10 minutes, and were coverslipped with Mowiol.

### Immunohistochemistry analysis

For quantification of cell type density, stainings of HLA-PLP, Iba1, CD3, CD20, CD79A, CD138, VWF, GFAP, and SOX10 were scanned (Axio slide scanner, x 20 magnification), and images were analyzed with Qupath ^55^. Based on the HLA-PLP staining the core, border zone, peri-lesional zone, and NAWM were annotated. Annotations were transferred onto the Iba1, CD3, CD20, CD79A, CD138, VWF, GFAP, and SOX10 scan, and the size was measured per annotation. Cell detection was performed on the nuclei staining, and positive cells were found with object classification. For each annotation, the number of microglia (Iba1), lymphocytes (CD3 + CD20), plasmablasts (CD79A + CD138), vascular cells (VWF), astrocytes (GFAP), and oligodendrocytes (SOX10) per mm^2^ were quantified with a cell profiler. The density of IGG^+^CD79A^+^ B-cells, FTH^+^HLA^+^ microglia, B2M^+^SOX10^+^ oligodendrocytes, ABCA2^+^SOX10^+^ oligodendrocytes, and QKI^+^SOX10^+^ oligodendrocytes was quantified by training a random trees classifier.

### Statistical analysis

Differences in proportions or density were tested with a quasi-Poisson general linear mixed model, with an offset of the total detected cells or area. Correlations were tested with Pearson’s correlation coefficient. All p-values were corrected for multiple testing with FDR. P-values were considered significant if <0.10, or if <0.05 for single-cell level zone comparison.

## Data availability

Data will be publicly available at the website of the Human Protein Atlas. Raw data will be available upon reasonable request.

