## Extended Fig. for "Microglial states determine lesion dynamics in multiple sclerosis"

**Extended data**

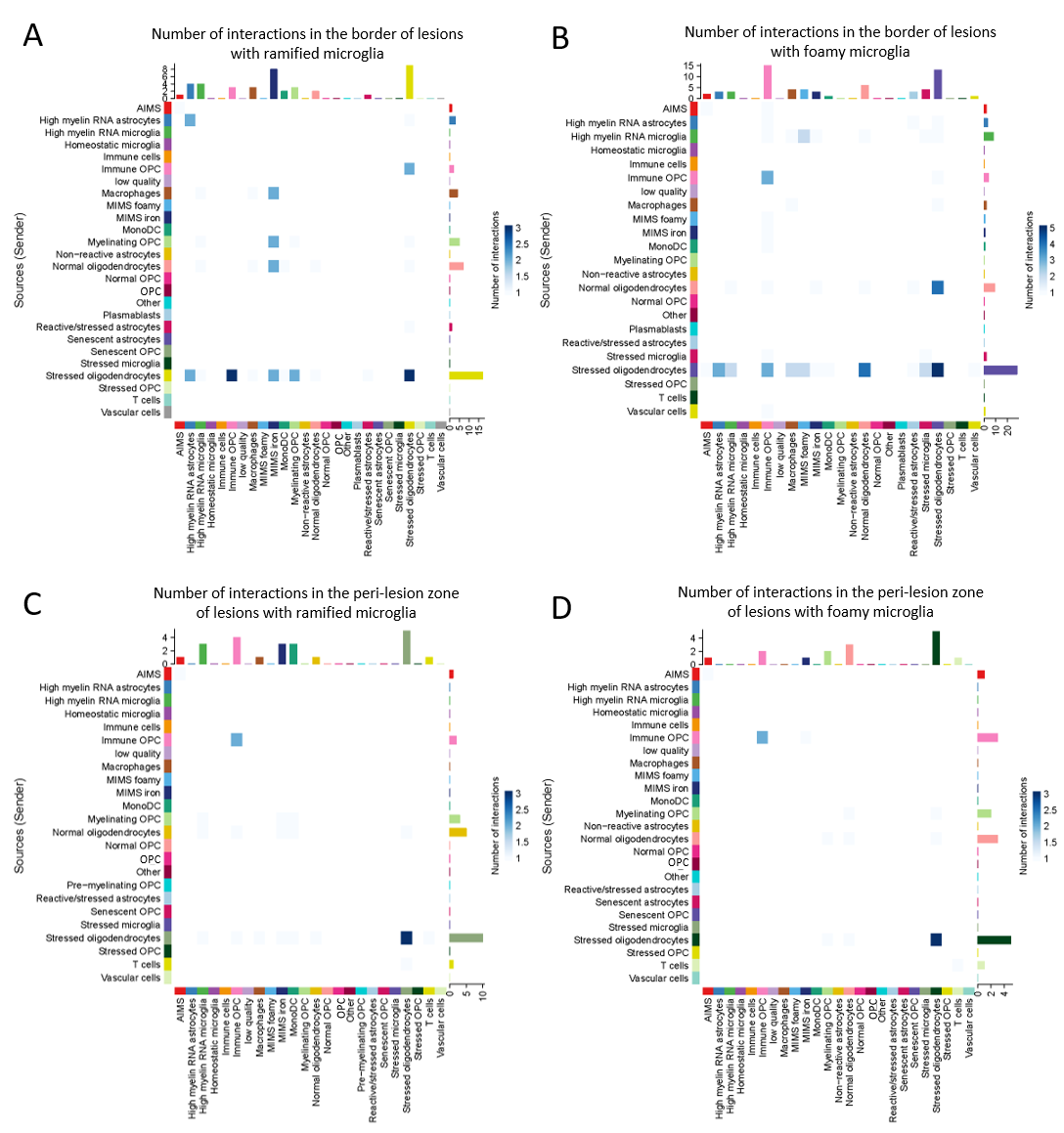

**Extended Fig. 1: Intercellular communication of cell types identified with CellChat**. A) Interactions in the border of mixed lesions with ramified microglia. B) Interactions in the border of mixed lesions with foamy microglia. C) Interactions in the peri-lesional zone of lesions with ramified microglia. D) Interactions in the peri-lesional zone of lesions with foamy microglia.

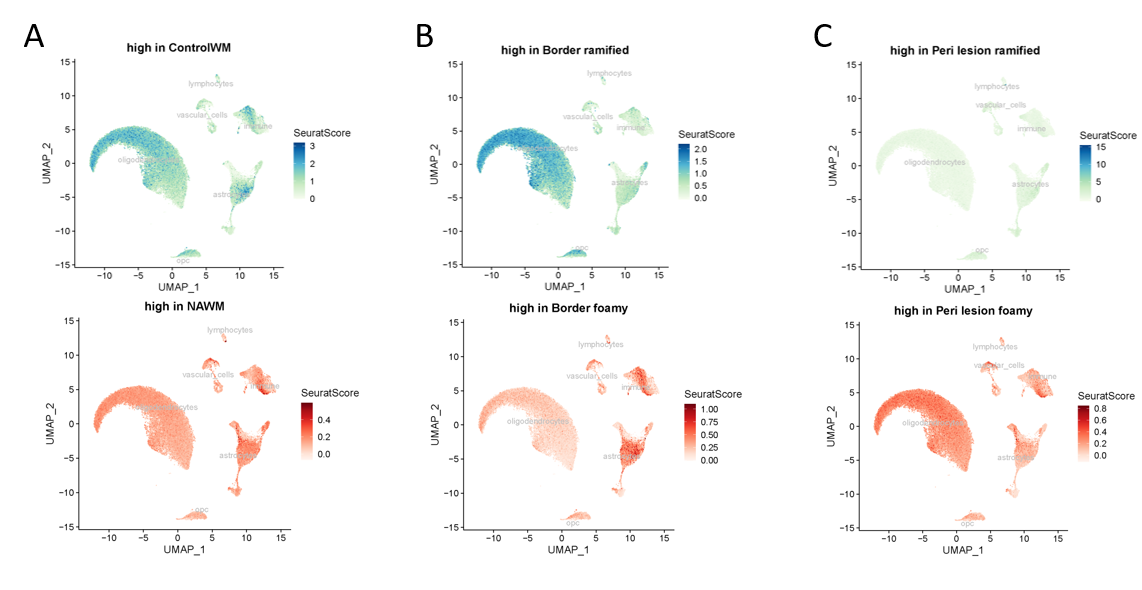

**Extended Fig. 2: Differentially expressed genes plotted onto the single-cell uMap of Absinta *et al.,* indicating major cell types affected**. A) De genes in NAWM compared to control WM. Genes with higher counts in the control WM (top panel) and with higher counts in the NAWM (lower panel). B) The border of mixed lesions with foamy microglia compared to the border of those with ramified microglia. Genes with higher counts in the border of mixed lesions with ramified microglia (top panel) and with higher counts in the border of those with foamy microglia (lower panel). C) The peri-lesional zone of mixed lesions with foamy microglia compared to the peri-lesional zone of those with ramified microglia. Genes with higher counts in the peri-lesional zone of mixed lesion with ramified microglia (top panel) and with higher counts in the peri-lesional zone of mixed lesions with foamy microglia (lower panel).

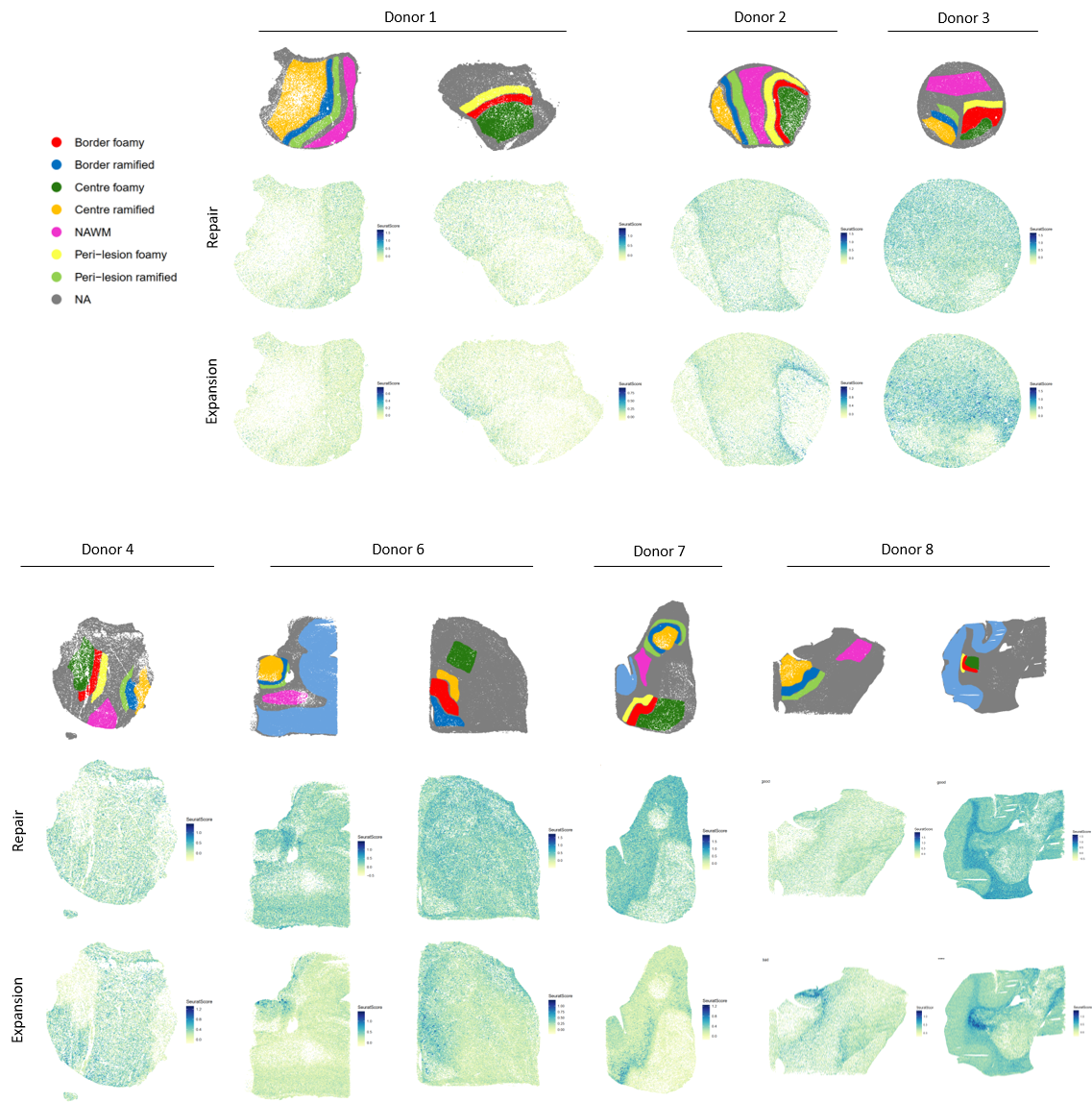

**Extended Fig. 3: Gene modules associated with repair or with expansion of lesions plotted onto the chip.** For each chip, zones of interest are indicated. Genes associated with repair were *APOD, CTNNB1, QKI, SBF2, BOK, CTTN, SHTN1, BCAS1, APOD, SHANK2, CDH4, SKI, NEAT1, DDR1*, those associated with expansion were *APOC1, APOE, APP, B2M, C1QA, C1QB, C1QC, C3, CD44, CD68, CD74, CD99, CHI3L1, CHIRL2, CHIT1, CLU, FABP5, FCGR1A, FCGR2A, FCGR2B, FTH1, FTL, GPNMB, HLA-A, HLA-DQB1, HLA-DRB1, IAH1, IFNGR1, IHA1, IGHG1, IGHG3, IGHM, IGLC2, IGLC3, IGLL5, JCHAIN, LIPA, MSR1, NCAM1, OLR1, DERPINA3, SPP1, UBB*.
